# Fluffy feathers: how neoptile feathers contribute to camouflage in precocial chicks

**DOI:** 10.1101/2020.07.26.221671

**Authors:** Veronika A. Rohr, Tamara Volkmer, Dirk Metzler, Clemens Küpper

## Abstract

Camouflage is a widespread strategy to increase survival. The plumage of precocial chicks often contains elements of disruptive colouration and background matching to enhance concealment. Chick plumage also features fringed feathers as appendages that may contribute to camouflage. Here, we examine whether and how neoptile feathers conceal the outline of chicks. We first conducted a digital experiment to test two potential mechanisms for outline diffusion through appendages: 1) edge intensity reduction and 2) luminance transition. Local Edge Intensity Analysis (LEIA) showed that appendages decreased edge intensity and a mean luminance comparison revealed that the appendages created an intermediate transition zone to conceal the object’s outline. The outline was most diffused through an intermediate number of interspersed thin appendages. Increased appendage thickness resulted in fewer appendages improving camouflage, whereas increased transparency required more appendages for best concealment. For edge intensity, the outline diffusion was strongest for a vision system with low spatial acuity, which is characteristic of many mammalian predators. We then analysed photographs of young snowy plover (*Charadrius nivosus*) chicks to examine whether neoptile feathers increase outline concealment in a natural setting. Consistent with better camouflage, the outline of digitally cropped chicks with protruding feathers showed lower edge intensities than the outline of chicks cropped without those feathers. However, the observed mean luminance changes were not consistent with better concealment. Taken together, our results suggest that thin skin appendages such as neoptile feathers improve camouflage. As skin appendages are widespread, this mechanism may apply to a large variety of organisms.

## Introduction

Avoiding detection either for protection from predators or to go unnoticed by potential prey is essential for individual survival. The threat of predation has led to the evolution of various camouflage mechanisms, which make potential prey more difficult to detect or recognize. The most prominent mechanism is visual camouflage that includes highly adaptive colouration strategies among animals (Stevens and Merilaita 2009). One strategy to achieve visual camouflage is background matching (also termed “crypsis” by Endler (1981)). For background matching, animals try to match colour, luminance and pattern of their background.

While background matching is one of the most common and frequently studied strategies of visual camouflage (Cott 1940, Endler 1981, Farkas et al. 2013, Allen et al. 2015, Stevens et al. 2017), another important mechanism is concealing the outline of the body. Thayer (1909) proposed that detecting the outline of their prey is one of the ways predators locate and identify their prey. In general, the detection of edges is an essential task for object recognition (Marr 1976, Tovée 1996). In this regard, disruptive colouration makes animals less detectable. It involves a set of markings that creates false edges within the animal hindering the detection or recognition of its true outline and shape or parts of it (Thayer 1909, Cott 1940, Stevens and Merilaita 2009). Cott (1940) suggested that structural modifications of the organism’s outline themselves could contribute to camouflage by creating an ‘irregular marginal form’. This makes the animal’s true body outline effectively diffused and hence makes it harder to detect (Cott 1940). Recently, support for the ‘irregular form’ hypothesis was found in an experimental study showing that false holes markings reduce avian predations (Costello et al. 2020).

Birds with their typically advanced vision and high plumage diversity have been featured prominently in camouflage research, either as predators or as prey (Cuthill et al. 2005, Skelhorn et al. 2010, Farkas et al. 2013, Lovell et al. 2013, Stevens et al. 2017, Pike 2018, Costello et al. 2020). When studying camouflage as an anti-predator defence in birds, much research has examined the clutches/eggs of ground-nesting birds (Stoddard et al. 2011, Ekanayake et al. 2015, Stevens et al. 2017). These studies revealed that ground-nesting birds may increase background matching through adaptive egg colouration that matches the nest site (Lovell et al. 2013, Stevens et al. 2017) and some species even improve the background matching of their clutches, by soiling their eggs to conceal them better (Mayani-Parás et al. 2015), using egg-matching nest materials (Gómez et al. 2018) or covering the clutch with debris or soil when predators approach (Troscianko et al. 2016).

However, not only eggs are vulnerable to predation. Chicks are also often targeted by predators. Precocial chicks leave their nest within a few hours of hatching. Initially, those chicks suffer from high mortality as they are limited in their mobility and hence highly vulnerable to predation (Colwell et al. 2007, Brudney et al. 2013, Eberhart-Phillips et al. 2018). To improve their survival, chicks rely on camouflage provided by their feathers especially during the first days of their lives. The plumage colouration of precocial chicks featured prominently in the description of camouflage mechanisms such as disruptive colouration (Thayer 1909, Cott 1940, Hill and McGraw 2006). However, we know surprisingly little about plumage characteristics that improve camouflage in chicks. Precocial chicks hatch fully covered with neoptile down feathers (Foth 2011). With maturation, the neoptile feathers are shed, and the natal plumage is replaced by the teleoptile feathers, which can be categorised into, e.g. flight, contour and down feathers (Stettenheim 1976). One striking feature of neoptile feathers is that they are protruding from the chick’s body. The unequal length of the very thin feathers creates a fringed feather region that may conceal the chick outline and hence make it harder to detect by predators.

In this study, we investigated whether neoptile down feathers improve camouflage through outline diffusion. Cott (1940) discussed this strategy of an ‘irregular marginal form’ mainly with examples of masquerade, where the irregular shapes of animals resemble elements of their environment, e.g. parts of plants. In contrast, we hypothesized that the fringed feathery outline helps the chick to better blend with the background by reducing edge contrasts and/or creating a transition zone of intermediate luminance.

In a first experiment, we explored the mechanism of outline diffusion by appendages in principle modelling a circular object with varying protruding appendages. We then used the Local Edge Intensity Analysis (LEIA) (van den Berg et al. 2019) to investigate whether appendages decreased the edge contrast of the object’s outline. Additionally, we investigated how appendage characteristics such as their density, thickness, transparency, and variation in background complexity and spatial acuity of the predator’s visual system affected edge intensity in the contour region. As a second mechanism, we tested whether appendages altered the luminance of a narrow ‘transition zone’ between object and background. We hypothesized that an intermediate mean luminance in the transition zone that reduces the contrast would help to blend the object better with the background.

In a second experiment, we tested whether the neoptile feathers contribute to the camouflage of precocial chicks. We analysed images taken from precocial snowy plover (*Charadrius nivosus*) chicks in natural habitats. Very young plover chicks rely on their crypsis to evade predation as they stay motionless on the ground when a threat is approaching (Colwell et al. 2007). We digitally cropped all chicks once with and once without protruding feathers and transferred them on to images of their hiding background taken after gently removing the chicks. For chicks cropped with their protruding feathers, we predicted the edge intensity of the chick outline to be reduced and the mean luminance difference of the transitions zone to be closer to intermediate optimum than for the images of those chicks cropped without their feathers.

## Material and methods

### Experiment 1: Proof of principle

As a proof of principle, we designed the first experiment to test whether appendages may help to conceal the outline. We created an image of a uniformly light grey coloured circular object with a size of 2950 pixels (px)/250.0 mm circumference and 470 px/39.8 mm radius on a dark grey background. The initial setup started with no appendages added to the outline (Figure 1a, ‘0’). We then added object-coloured appendages (i.e. lines of 1 Pt/4 px/0.4 mm thickness and 118 px/10.0 mm length) with regular intervals resembling protruding neoptile chick feathers orthogonally to the object outline (‘Basic Scenario’, Figure 1a). The first image with appendages had 32 appendages added to the outline (Figure 1a, ‘32’). We then doubled the number of appendages stepwise creating denser spaced appendages to the outline until the extended outline was completely filled (Figure 1a, ‘full circle’). For the vision of a simulated predator, we used the spatial acuity from humans (*Homo sapiens*, 72 cycles per degree, cpd) (Land 1981; Land and Nilsson 2012; Caves and Johnsen 2018) in the basic scenario. The full details for the parameters are provided in Table S1 (a to g).

**Figure 1:**
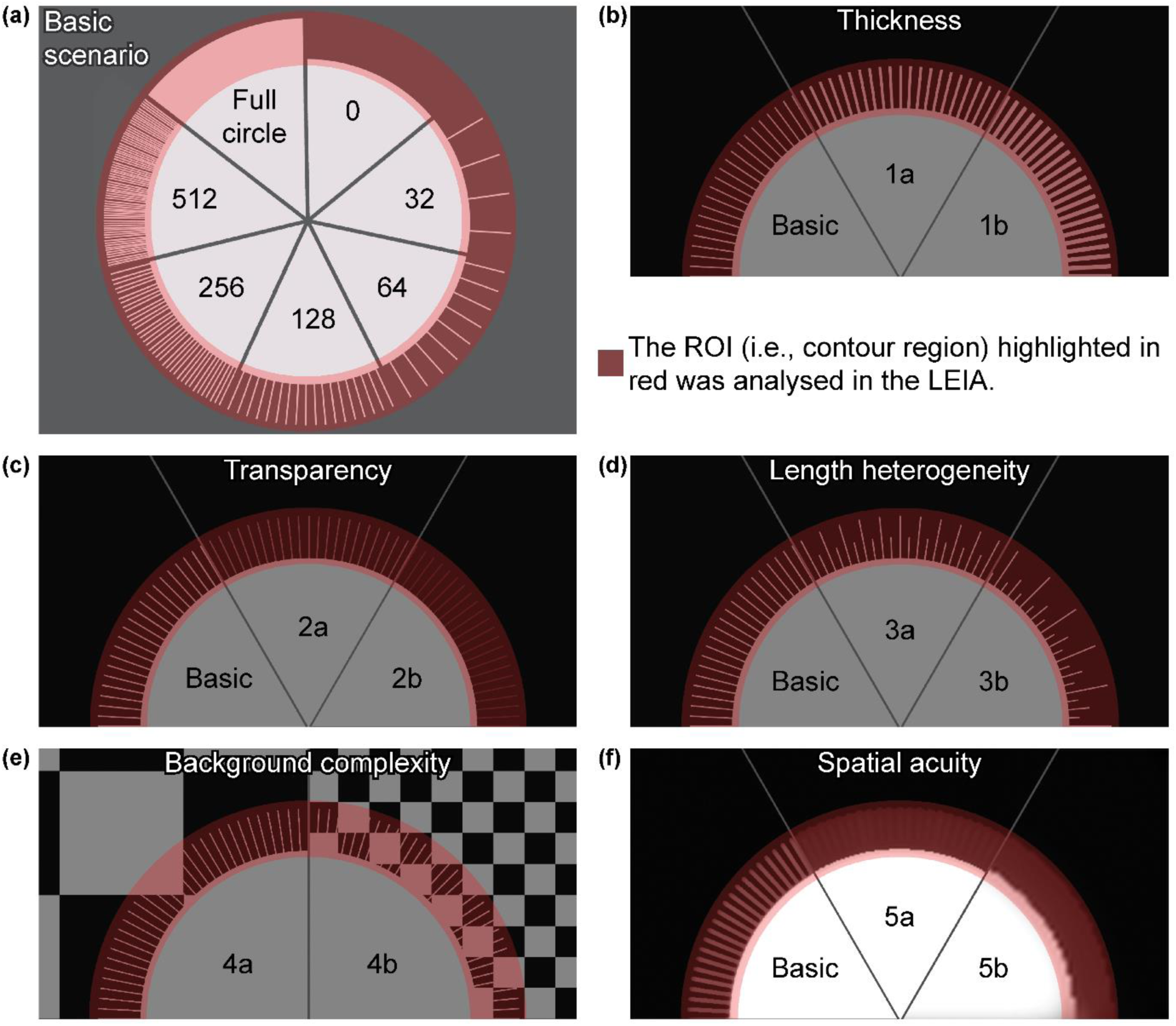
(a) Basic Scenario: Seven stages of the artificial chick setup with varying number of thin, non-transparent appendages having all the same length. (b) Scenario 1: varying appendage thickness applied to the Basic Scenario. (c) Scenario 2: varying appendage transparency applied to the Basic Scenario. (d) Scenario 3: varying appendage length heterogeneity applied to the Basic Scenario. (e) Scenario 4: varying background complexity with artificial chessboard backgrounds. (f) Scenario 5: high, medium and low visual acuity applied to the Basic Scenario. (a – f) The analysed region of interest (ROI) is highlighted in red for clarification only.

To further explore the mechanism, we altered appendage characteristics, background and the spatial acuity of the predator. First, we increased appendage thickness to 2 Pt/8 pixels/ 0.7 mm (Scenario 1a) and 3 Pt/12 pixels/1.1 mm (Scenario 1b) resulting in decreased inter-appendage intervals (Figure 1b and Table S1, h to u). Second, we changed appendage transparency to 25 % (Scenario 2a) and 50 % transparency (Scenario 2b) (Figure 1c and Table S1, v to ai). Third, we varied the appendage length heterogeneity; half of the appendages having 50 % of the length (Scenario 3a), and half of the appendages at 25 % and one quarter at 50 % of the original appendage length (Scenario 3b) (Figure 1d and Table S1, aj to aw). Fourth, we investigated the effect of background complexity on the detectability of the outline. As background, we used a chessboard pattern with large squares (346 pixels/29.3 mm, Scenario 4a) and with small squares (86 pixels/7.3 mm, Scenario 4b) (Figure 1e and Table S1, ax to bk). Fifth, we altered the spatial acuity to test whether or how the visual systems of different predators would affect detectability. We simulated the spatial acuity of a corvid predator (30 cpd, Scenario 5a) and canid predator (10 cpd, Scenario 5b) (Figure 1f and Table S1, bl to by), the two most common predators of ground-nesting plovers (Burrell and Colwell 2012, Ekanayake et al. 2015, Ellis et al. 2020). This range also covered other potential predators (Table S2).

We did not account for differences in colour vision between different predators as the setup mostly consists of greyscale images that predominantly differ in luminance. Note that in many animals, visual acuity is greater for achromatic than chromatic stimuli (Giurfa et al. 1997, Endler et al. 2018).

We conducted visual modelling and visual analysis using the Quantitative Colour Pattern Analysis (QCPA) framework (van den Berg et al. 2019) integrated into the Multispectral Image Analysis and Calibration (MICA) toolbox (Troscianko and Stevens 2015) for ImageJ version 1.52a (Schneider et al. 2012). We converted the generated images into multispectral images containing the red, green and blue channel in a stack and transformed them further into 32-bits/channel cone-catch images based on the human visual system, which are required by the framework. To create the luminance channel, we averaged the long and medium wave channel, which is thought to be representative of human vision (Livingstone and Hubel 1988) (Figure S1a). We modelled the spatial acuity with Gaussian Acuity Control at a viewing distance of 1300 mm and a minimum resolvable angle (MRA) of 0.01389 (Figure S1b). To increase biological accuracy, we applied a Receptor Noise Limited (RNL) filter that reduces noise and reconstructs edges in the image. The RNL filter used the Weber fractions “Human 0.05” provided by the framework (longwave 0.05, mediumwave 0.07071, shortwave 0.1657), luminance 0.1, 5 iterations, a radius of 5 pixels and a falloff of 3 pixels (Figure S1c) as specified in van den Berg et al. (2019).

#### Local Edge Intensity Analysis

To test for the detectability of the outline, we used LEIA (van den Berg et al. 2019), which is conceptually similar to the boundary strength analysis (Endler et al. 2018). Boundary strength analysis requires an image with clearly delineated (clustered) colour and luminance pattern elements. However, a large degree of subthreshold details, which may be still perceived by the viewer gets lost in the clustering process. LEIA has the advantage of not requiring such a clustered input and therefore can be directly applied to RNL filtered images. LEIA measures the edge intensity (i.e. the luminance contrast) locally at each position in the image. The output image displays ΔS values in a 32-bit stack of four slices, where each slice shows the values measured in different angles (horizontal, vertical and the two diagonals, for more details, see van den Berg et al. (2019)).

We ran LEIA on the chosen region of interest (ROI) with the same Weber fractions used for the RNL filter. The ROI was the contour region, a 180 pixel-wide band that included the area of the appendages extended by 30 pixels towards the object inside and towards the outside (Figure 1a). We log-transformed the ΔS values as recommended for natural scenes (Troscianko and van den Berg 2020) to make the results comparable to the natural background images used in Experiment 2 (see below). To test whether the size of the ROI affected our results, we ran an additional analysis using a 1500 × 1500 pixel-wide rectangle surrounding the object as the ROI, which included a bigger area of the background and the full object inside (Figure S2a).

We extracted the luminance ΔS values from the four slices of the output image stack in ImageJ and stored them in separate matrices for further analysis using R version 3.5.3 (R Core Team 2019). ImageJ generally assigned values outside the chosen ROI to zero. Thus, we first discarded all values of zero. We then set all negative values that arose as artefacts in areas without any edges to zero, in order to make them biologically meaningful. We then identified the parallel maximum (R function *pmax ()*) of the four interrelated direction matrices and transferred this value to a new matrix.

High luminance and colour contrasts imply high conspicuousness (Endler et al. 2018). Consequently, a lower luminance contrast leads to lower conspicuousness and therefore, better camouflage. As the outline is an important cue for predators locating and identifying a prey item (Thayer 1909), we assumed that especially low contrasts in the outline of an object improve camouflage. Thus, a reduction of edge intensity in the object outline by the appendages indicates a camouflage improvement. To test whether the object outline became less detectable we compared the edge intensity of the outline pixels in the basic scenario without appendages (Table S1, a) with corresponding pixels from other scenarios. The outline pixels were characterised by high edge intensity and constituted a prominent peak. They comprised 1.59 % of all pixels in the analysis focused on the contour region (see Results, Figure 2a). For all scenarios, we calculated the mean edge intensity of the high edge intensity pixels (HEI pixels) and identified the changes with parameter variation.

**Figure 2:**
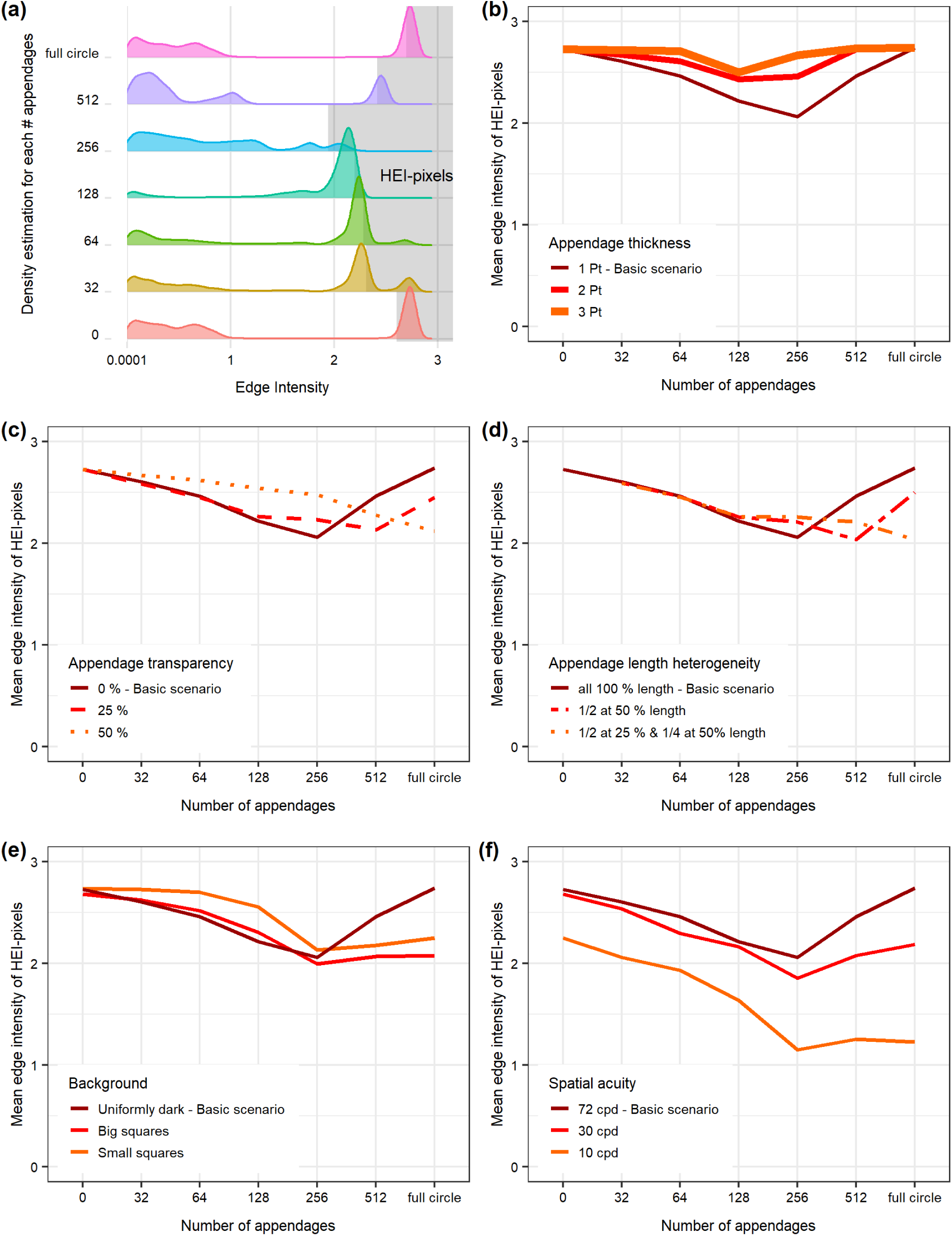
Local edge intensity analysis of the contour region in the artificial object experiment. (a) Ridgeline plots showing the density distribution of the edge intensity according to number of appendages. The highest 1.59 % of the pixels are shaded in grey (High edge intensity pixels, HEI pixels). (b) Scenario 1: variation in appendage thickness. (c) Scenario 2: variation in appendage transparency. (d) Scenario 3: variation in appendage length. (e) Scenario 4: variation in background complexity. (f) Scenario 5: variation in spatial acuity.

#### Mean Luminance Comparison

For the Mean Luminance Comparison (MLC), we analysed the same images as with the LEIA. We divided the filtered image into three regions of interest (ROIs) (Figure 3a). 1) The object region included the whole object inside up to 20 pixels next to the object outline. 2) The appendage region was an 80 pixel-wide band including only the area covered by appendages. It started 20 pixels outside the object outline and reached up to 20 pixels before the boundary created by the appendages (appendage-boundary). 3) The background region ranged from 20 pixels outside the appendage-boundary to a 1500×1500 pixel-wide rectangle surrounding the object. A buffer zone of 40 pixels between all three regions was excluded from the analysis to ensure a clear separation of the regions. In the luminance channel of each image, we measured the mean luminance in the three regions and compared them subsequently. Luminance values range from 0 to 1.

**Figure 3:**
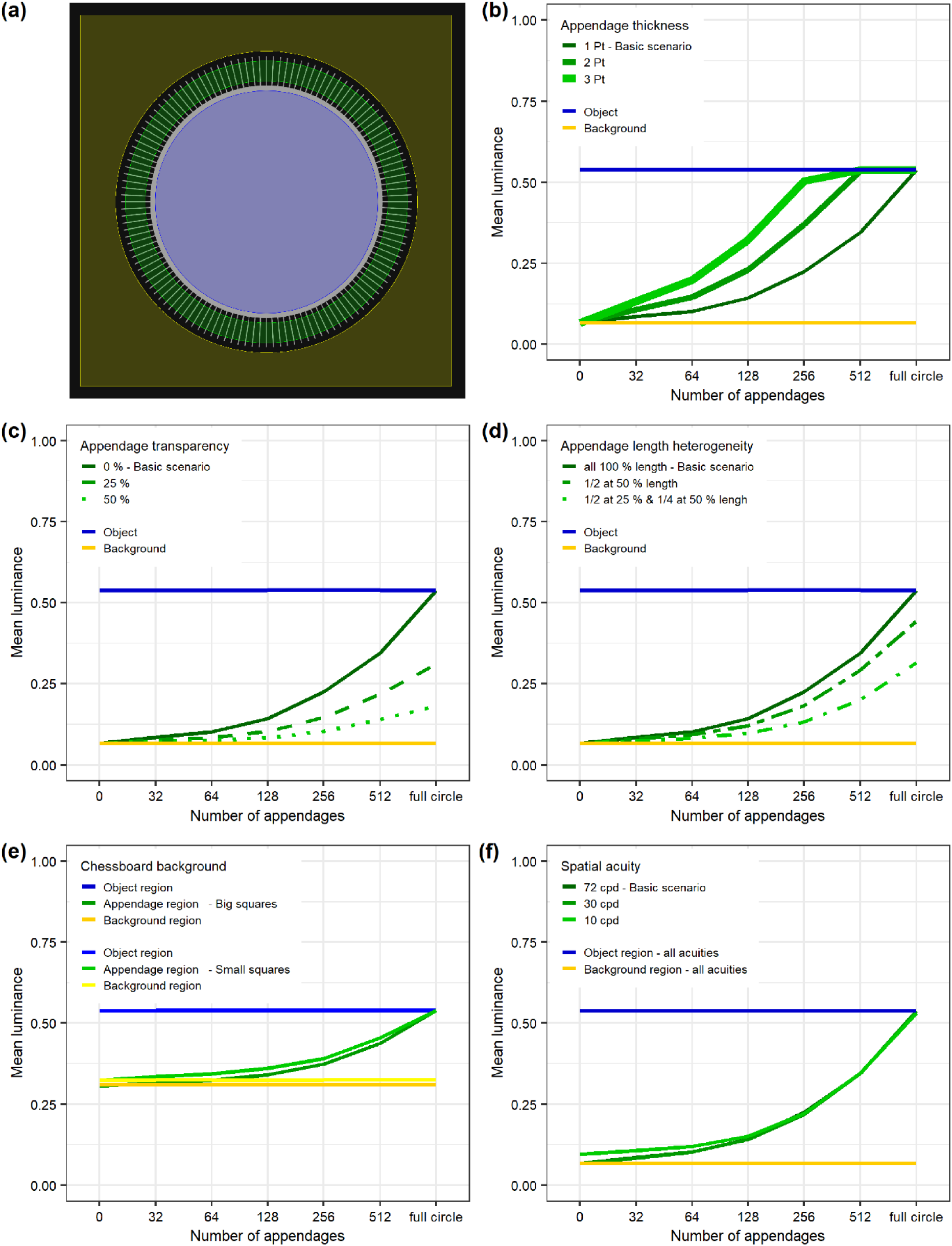
Mean Luminance Comparison in experiment 1. (a) The three regions of interest (ROIs) analysed were object region (blue), background region (yellow) and appendage region (green). The ROIs were separated by 40 pixels to ensure a clear separation of the regions. (b) Scenario 1: variation in appendage thickness. (c) Scenario 2: variation in appendage transparency. (d) Scenario 3: variation in appendage length. (e) Scenario 4: variation in background complexity. (f) Scenario 5: variation in spatial acuity. Note that the 30 and 72 cpd curves overlap fully.

According to background matching, objects that differ more in luminance from the background are more conspicuous and hence less well camouflaged (Endler 1981). We assumed that detectability based on possible luminance differences between object and background are weakened by the appendages as they form a transition zone helping to blend the object better into the background. Accordingly, from a camouflage perspective, the appendage region would provide an optimal transition zone when its mean luminance is exactly the mean of the object and background region’s luminance.

### Experiment 2: Chick photographs

Using pictures of young snowy plover chicks hiding when approached by a predator, we tested if protruding neoptile feathers helped to conceal the chicks’ outline and therefore improve their camouflage.

We studied snowy plovers in their natural environment at Bahía de Ceuta, Sinaloa, Mexico. The breeding site consists of salt flats that are sparsely vegetated and surrounded by mangroves (Cruz-López et al. 2017). General field methodology is provided elsewhere (Eberhart-Phillips et al. 2020). In 2017, we took photographs of young (one to three days old) chicks hiding on the ground, that had already left the nest scrape. To photograph the chicks, two observers approached free-roaming families with two mobile hides (Székely et al. 2008) within the period one hour after sunrise and one hour before sunset. At a distance of 100-200 m, one observer acted as ‘predator’, left the hide and openly approached the brood while the second observer kept watching the chicks. The chicks responded by crouching to the ground and staying motionless while the parents were alarming. The second observer directed the ‘predator’ to the approximate hiding place. When searching for the chicks, we took great care to reduce the number of steps to avoid modification of the ground through our tracks.

Once the first chick had been found, the second observer joined the predator and took chick photographs. We used a Nikon D7000 camera converted to full spectrum including the UV range (Optic Makario GmbH, Germany) and a Nikkor macro 105 mm lens that allows transmission of light at low wavebands. The equipment was chosen because calibration data were available for this combination (Troscianko and Stevens 2015). Each hiding background was photographed with and without the chick using a UV pass filter for the UV spectrum and a UV/IR blocking filter (“IR – Neutralisationsfilter NG”, Optic Makario GmbH, Germany) for the visible spectrum. The camera was set to an aperture of f/8, ISO 400 and the pictures were stored in “RAW” file format. We used exposure bracketing to produce three images to ensure that at least one picture was not over or underexposed. A 25 % reflectance standard (Zenith Polymer TM) placed in the corner of each picture enabled a subsequent standardizing of light conditions.

In total, we took pictures of 32 chicks from 15 families. For 21 chicks we obtained photographs suitable for further analyses with an unobstructed view to the entire chick and only one chick per photograph. Of these, we randomly selected pictures of 15 chicks. Unfortunately, it was not possible to obtain proper alignment of visual and UV pictures in ImageJ as either chick or camera moved slightly in the break between changing filters for the two settings. Therefore, we restricted our analyses to human colour vision and discarded the UV pictures for further analysis.

In each picture, we manually selected the chick outline and the feather-boundary as a basis for the ROIs (Figure 4a-c). The chick outline included bill, legs, rings and all areas densely covered by feathers without background shining through. We then marked the feather-boundary, i.e., the smoothened line created by the protruding neoptile feather tips. In the next step, we transferred images of chicks with or without protruding feathers, i.e. cropped at feather-boundary or chick outline, respectively, and inserted them into a uniform or the natural background. First, we cropped the chick without protruding feathers and transferred it into a uniform black background. Second, we cropped the chick including all feathers and inserted it into exactly the same hiding spot on the picture of the natural background (Figure 4b). Third, we cropped the chick excluding the protruding feathers and transferred it into the natural background (Figure 4c).

**Figure 4:**
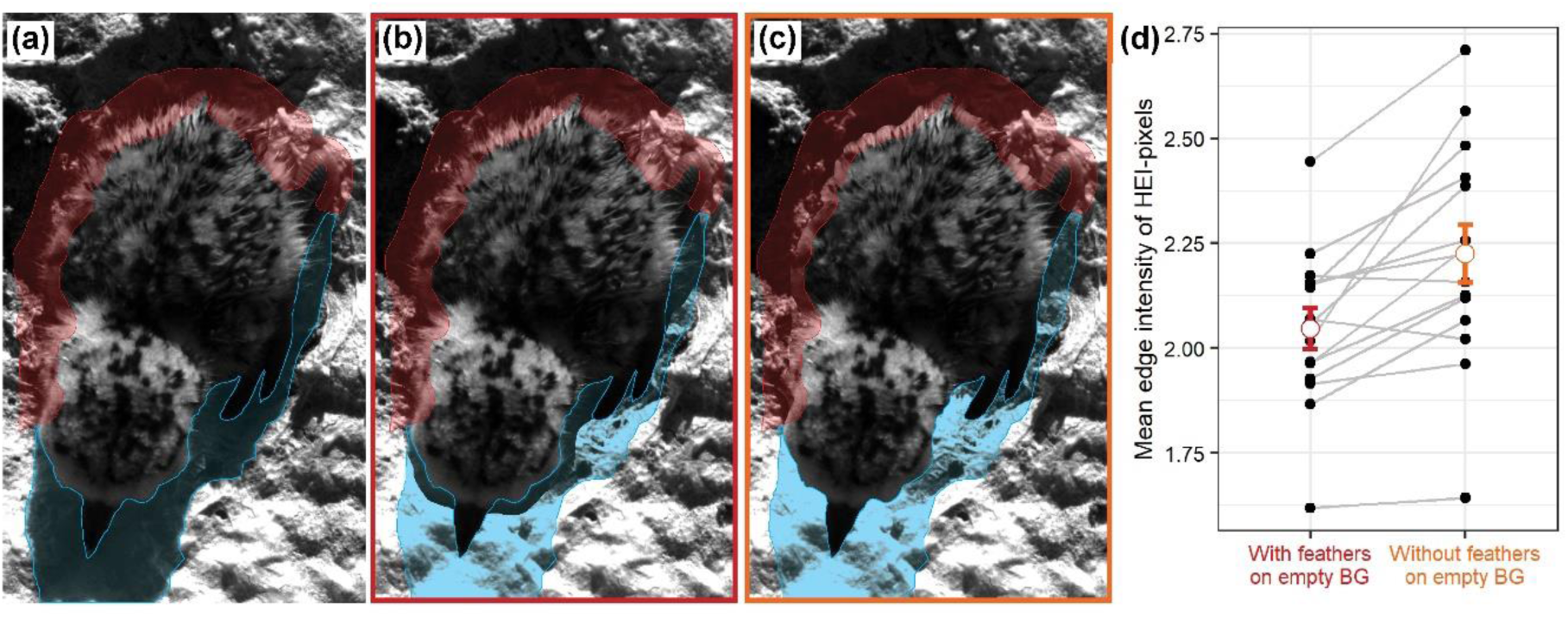
(a) A snowy plover chick hiding on the ground from an approaching predator (b) cropped chick transferred to image of empty natural background with neoptile contour feathers protruding the outline (c) cropped chick transferred without protruding neoptile contour feathers. The contour region (red) as the region of interest was analysed in the Local Edge Intensity Analysis. Areas, where the background was shaded by the chick in the original image (blue), were excluded from the analysis. (d) Mean edge intensity of the HEI pixels in the contour region with and without feathers for 15 snowy plover chicks (t = 4.365, df = 14, p-value < 0.001). Measurements are paired by chick ID. The error bars indicate group mean ± standard error.

#### Local Edge Intensity Analysis

We then proceeded with LEIA following the protocol of experiment 1 with the following changes. Again, the selected ROI was the contour region ranging from the chick outline extended by 30 pixels towards the chick inside to the feather-boundary extended by 30 pixels towards the outside. We excluded all areas of the ROI that showed a shadow of the chick as the chicks’ shadow was missing on the empty natural background images to which the cropped chicks were transferred to (Figure 4a-c). We used the images of the cropped chicks on the black background to determine the threshold of the HEI pixels according to the protocol of experiment 1 for each chick separately. For each cropped chick that was transferred to the picture with the natural background, we compared the mean edge intensity of the HEI pixels provided by LEIA with and without protruding feathers (Figure 4b-c) using a two-sided paired t-test.

#### Mean Luminance Comparison

We also calculated mean luminance differences for each chick using the same cropped photographs as for the LEIA. Similar to the artificial object experiment, the chick region included everything inside the chick outline, the background region included everything outside the feather-boundary up to a 1500×1500 pixel-wide rectangle surrounding the chick and the feather region (FR) was between chick outline and feather-boundary. Note that the FR is different from the contour region, which additionally includes a small part of chick and background region. We reduced the FR by excluding all areas that were shaded by the chick since this shadow was missing on the empty background images. Additionally, we excluded the buffer zone (Figure 3a, the area between the coloured regions) to cover the whole variation in feather density in the FR (Figure 5a-b). Close to the chick outline, the feathers were still relatively dense thinning more and more towards the feather-boundary as they were very variable in length.

**Figure 5:**
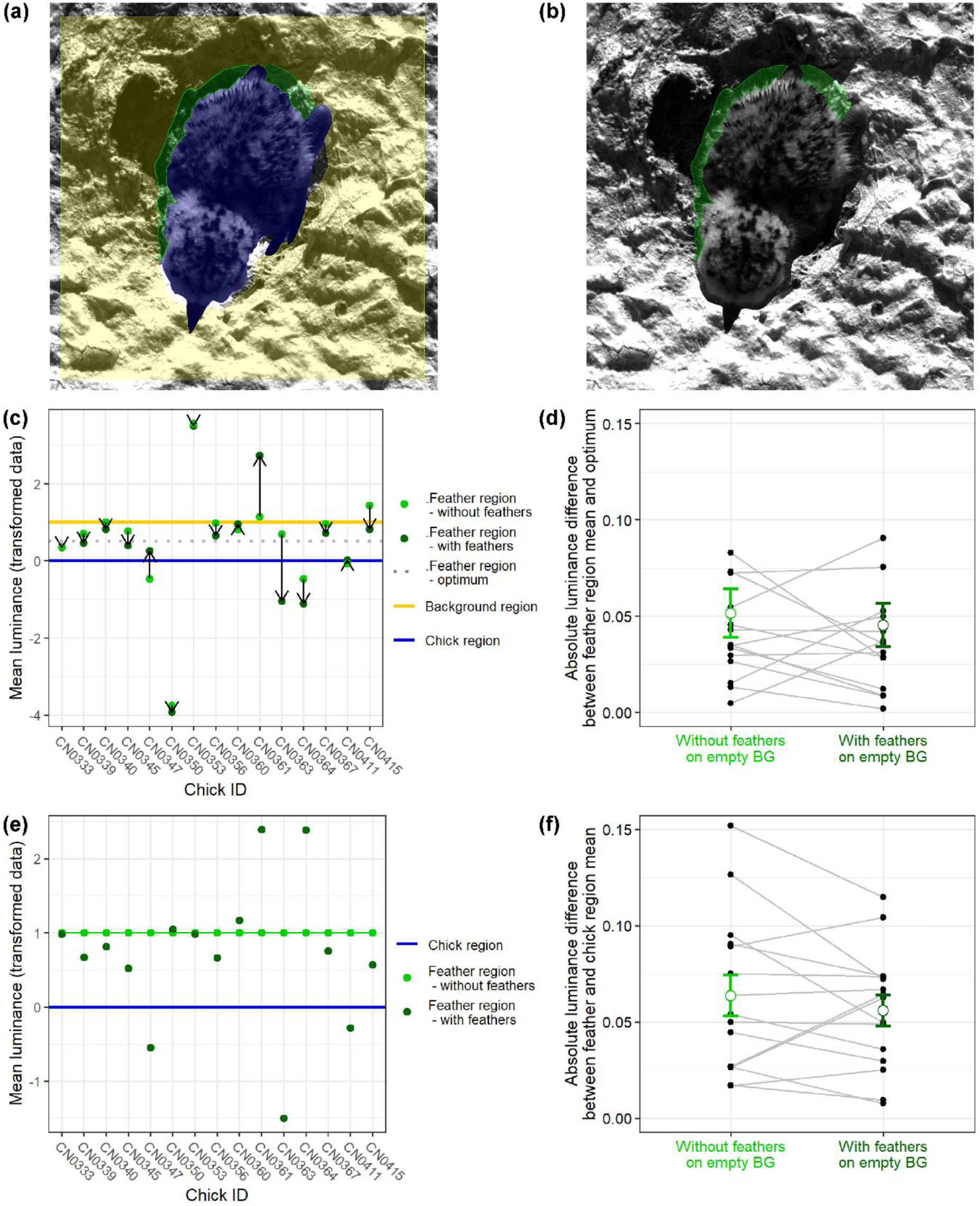
Mean Luminance Comparison in experiment 2. The regions of interest (ROIs) analysed were (a) chick (blue) background (yellow) and feather region (FR) without feathers (green) measured in scene 2 and (b) the FR with feathers (green) measured in scene 1. (c) Measurements were transformed so that the chick region (blue) was always “0” and the background region (yellow) “1”. The FR optimum (grey, dotted) portrays the mean of chick and background region. The arrows indicate the direction in which the value of the FR was shifted when the feathers were present. (d) Absolute luminance difference between FR mean and optimum with and without feathers.* (e) Measurements were transformed so that the chick region (blue) was always “0” and the FR without feathers (light green) “1”. (f) Absolute luminance difference between chick region and FR mean with and without feathers.* *Measurements are paired by chick ID. The error bars indicate group mean +/-standard error.

For each chick, we measured the mean luminance of all three regions in the luminance channel of the image containing the chick without feathers (Figure 5a). The FR we measured again in the image containing the chick with feathers (Figure 5b).

In theory, the best transition zone between chick and background that reduces the outline of the chick against the background the most should have an exactly intermediate luminance between chick and background region. In a first analysis, we checked whether the absolute distance of mean luminance of the FR with feathers was closer to those optimal values than without feathers. Because the luminance data were not normally distributed according to the Shapiro-Wilk normality test we conducted a Wilcoxon paired signed rank test. To compare the data graphically in an intuitive way, we transformed the values so that the chick region always was the reference with a value of 0, the background region became 1. The two values measured in the FR stayed in their initial relative distance to chick and background value.

The FR generally was quite narrow compared to chick and background region and its effect probably acts predominantly from close proximity. Therefore, we focussed the next analysis only on chick and FR. We assumed that the chick to a certain extent differs in luminance from its immediate background in the FR and that including the feathers decreases this difference and thus possibly improves the camouflage. Therefore, we compared the absolute distances between the mean luminance of chick region and FR with and without feathers. As the data were normally distributed according to the Shapiro-Wilk normality test we conducted a two-sided paired t-test.

For an easier comparison of the measurements, we transformed the luminance values in this analysis. The chick region again was the reference with a value of 0. As the background region was excluded, we scaled the FR without feathers to 1. The value measured in the FR with feathers stayed in its initial relative distance to the other two values.

The analysis aimed to check if the FR meets the basic requirement of a transition zone having intermediate luminance. Thus, we checked whether the mean luminance value of the FR with feathers fell between the one of chick region (mean luminance = 0) and FR without feathers (mean luminance = 1) constituting the immediate surrounding background to account for the local scale. We calculated the probability for the FR with feathers of having a value between 0 and 1 when randomly distributed. For this, we drew a random sample (n = 10,000) from a normal distribution with the mean and standard deviation in the transformed data. Then, we ran an exact binomial test to determine whether the observed intermediate luminance value was different from the expected value.

## Results

### Experiment 1: Artificial object

#### Local Edge Intensity Analysis

All images showed multimodal density distributions of pixels (Figure 2a). Pixels showing the highest edge intensities were found at the object outline. These HEI pixels showed prominent modal peaks in all multimodal density distributions (Figure 2a). For the object without appendages, 1.59% of pixels made up the distinct modal area with a mean edge intensity of 2.7 (Figure 2a, ‘0’). Consequently, we used a threshold of 1.59% to define HEI pixels for all images. Adding appendages reduced the mean edge intensities of the HEI pixels with the lowest mean edge intensity reached in the image with 256 appendages (Figure 2a-b).

##### Appendage characteristics

Increasing appendage thickness (Scenario 1) resulted in overall higher mean edge intensities suggesting higher detectability than in the basic scenario. With thicker appendages, the lowest mean edge intensity of the HEI pixels was reached already with 128 appendages. Images with more than 128 appendages had higher mean edge intensity values implying a deterioration of camouflage (Figure 2b). Increasing appendage transparency (Scenario 2) yielded overall slightly higher mean edge intensities than observed in the basic scenario. The lowest mean edge intensities were reached with more appendages than in the basic scenario (Figure 2c) with the minimum mean edge intensity shown for 512 appendages at 25 % transparency and the full circle of appendages at 50 % transparency (Figure 2c). Increasing appendage length heterogeneity (Scenario 3) yielded the same low mean edge intensity values as the basic scenario (Figure 2d). However, more appendages were required to reach minimal mean edge intensity values than in the basic scenario. The minimum mean edge intensity was reached with 512 appendages when half of the appendages had 50 % of the length or with the full circle when half of the appendages had 25 % and a quarter had 50 % of the length (Figure 2d).

##### Background complexity and spatial acuity

Introducing background complexity (Scenario 4) resulted in similar mean edge intensities of the HEI pixels for 256 appendages as in the basic scenario for large squares. The ROI on the background with small squares showed slightly higher mean edge intensities for the HEI pixels than for the background with large squares. More appendages did not lead to such a pronounced increase of mean edge intensities as in the basic scenario (Figure 2e). Lowering the spatial acuity of the perceiver (Scenario 5) decreased the mean edge intensity severely. At a spatial acuity of 10 cpd, the minimum mean edge intensity of the HEI pixels in the image with 256 appendages was only half of the value obtained in the basic scenario (Figure 2f).

##### ROI Size

Changing the ROI size and examining a larger part of background and object (Figure S2a) produced qualitatively similar results (Figure S2b-d, f) except for variation in background complexity (Scenario 4). In that scenario, the number of appendages had no influence on the mean edge intensity of the HEI pixels (Figure S2e) for the enlarged ROI.

#### Mean Luminance Comparison

The mean luminance of the area covered by appendages (appendage region) was generally intermediate between the luminance of object and background across all scenarios indicating the formation of a luminance transition zone (Figure 3b-f). Without appendages, the appendage region’s mean luminance was the same as the one of the background region. With an increasing number of appendages, the appendage region’s mean luminance became more and more similar to the object region’s luminance until they were identical when the appendages formed a full circle (Figure 3b, dark green curve).

##### Appendage characteristics

Increasing the appendage thickness in Scenario 1 caused the appendage region’s luminance to converge sooner with the object region’s luminance. The optimum was also reached sooner, between 128 and 256 appendages at 2 Pt thickness and around 128 appendages at 3 Pt thickness respectively (Figure 3b). Having 128 appendages of 3 Pt thickness was the best parameter combination tested. In this image, approximately 50 % of the appendage region’s area was covered with appendages. This suggests for the basic scenario that the optimal intermediate luminance would have been reached for objects that have between 256 and 512 appendages (Figure 3b, ‘1Pt’), when 50 % of the appendage region would have been covered by appendages. In contrast, with increasing appendage transparency (Scenario 2) more appendages were needed to reach the same luminance values compared to the Basic Scenario. At 25 % transparency, the full circle of appendages was needed to reach the optimum intermediate value and with 50% transparency, the intermediate value could not be reached at all (Figure 3c). Similarly, with increasing appendage length heterogeneity (Scenario 3) more appendages were required to reach the optimum but it was obtained when half of the appendages had 50 % of the length as well as when half of the appendages had 25 % and a quarter had 50 % of the length (Figure 3d).

##### Background complexity and spatial acuity

Increasing the background complexity did not affect the curve trajectories in the transition zone. Without appendages, the appendage region’s mean luminance was similar to the background’s luminance and became increasingly similar to the object’s luminance when raising the number of appendages until they converged with a full circle of appendages (Figure 3e). Likewise, lowering the spatial acuity in Scenario 4 did not clearly change the curve trajectory in the transition zone (Figure 3f).

### Experiment 2: Chick photographs

#### Local Edge Intensity Analysis

For eight of the 15 analysed chicks, the empty background image was slightly shifted because of a camera movement. Therefore, we corrected their position manually to place the chicks exactly at the same spot in the empty background.

After removing the areas of the ROIs where the chick shaded the background, we were able to analyse on average 72 % of the contour region with LEIA. Across the ROIs of the 15 chicks, the mean threshold for the HEI pixels was 0.9826 (Table S3). Consequently, we compared on average 1.74 % of the pixels between photographs of cropped chicks with and without the protruding neoptile feathers.

For 13 of 15 chicks (87 %), the mean edge intensities of HEI pixels were lower for the cropped image of each chick with protruding neoptile feathers (e.g. Figure 4b) than for the corresponding images without protruding neoptile feathers (e.g. Figure 4c). Accordingly, images including the protruding feathers showed lower mean edge intensities of HEI pixels than those excluding them (Figure 4d, paired t-test: t = 4.365, df = 14, p-value < 0.001). The mean edge intensity difference of HEI pixels between measurements with and without feathers was 0.178 (95 %CI: 0.091, 0.265).

#### Mean Luminance Comparison

For the MLC, presence of protruding neoptile feathers did not contribute to creating a transition zone between chick and background, as we did not observe more intermediate mean luminance values in the ROI in comparison to ROI of chick pictures without protruding feathers (Figure 5c-f).

After removing the areas that were shaded by the chicks, on average 73 % of the FR remained for the MLC (Table S3). This value was slightly different from the 72 % that remained of the contour region in the LEIA because FR and contour region differed in size and the extent to which they were shaded by the chick.

Presence of feathers did not change the mean luminance of the transition zone adaptively. The distance of the mean luminance of the FR to the optimal intermediate value was in 9 of 15 chicks (60 %) shorter with than without feathers (Figure 5c-d). The Wilcoxon paired signed rank test showed no clear difference between the distribution of the two groups (p = 0.45).

Feathers did not make mean luminance of the FR more similar to the mean luminance of the chick region. Although the distance of the mean luminance of the FR with feathers to the mean luminance of the chick region was in 10 of 15 cases (66.7 %) shorter than without feathers (Figure 5f), there was no clear difference between images with and without protruding feathers (t = 1.1263, df = 14, p = 0.28). The mean luminance difference between the measurements with and without feathers was 0.0077 (95 %CI: −0.0070, 0.0224).

The FR with feathers had an intermediate mean luminance between the chick region and the FR without feathers in 8 of 15 chicks (53 %) (Figure 5e). In the random sample (n = 10,000) from a normal distribution with the same mean and standard deviation as observed in the transformed data, 38 % of the values were intermediate between 0 and 1. Including the protruding feathers, we observed a proportion of 0.53 (95 %CI: 0.27, 0.79) intermediate values, however, this was not clearly different from expected by chance (p = 0.29).

## Discussion

The plumage of newly hatched chicks has several known functions. The feathers are important for thermoregulation (Wekstein and Zolman 1971). Plumage colour variation is also an important signal that may reveal chick condition and facilitate individual recognition for parents (Johnsen et al. 2003, Hill and McGraw 2006, Lyon and Shizuka 2020). In precocial chicks, the plumage provides camouflage through cryptic colouration (Cott 1940, Hill and McGraw 2006). Here we tested whether neoptile feathers help to conceal the outline of chicks to make them harder to detect for predators. Our results from a proof of principle analysis (experiment 1) and analysis of real chick images in their natural environment (experiment 2) suggest that appendages, such as protruding neoptile feathers, improve concealment of the object outline, particularly by decreasing the edge intensity. Weak contrast edges are associated with low conspicuousness (Endler et al. 2018). This enhances diffusion of the outline and decreases detectability as the shape is an important cue for predators locating and identifying a prey item (Thayer 1909).

In the artificial setup (experiment 1), appendages both reduced edge intensity and created a transition zone with an intermediate mean luminance in the appendage region suggesting that both mechanisms help to conceal the object outline. However, when analysing the impact of neoptile feathers on outline concealment of chicks in their natural background (experiment 2), we found that the presence/absence of protruding feathers did only change edge intensity but not mean luminance of the ROI in the predicted way. ROIs on images where the chick was cropped including its protruding feathers had lower edge intensity but no consistent change in the intermediate luminance was found. This suggests that the lowering of edge intensity, which we analysed through LEIA (van den Berg et al. 2019) is a better mechanism for outline diffusion than creating a transition zone with intermediate luminance for concealing the outline of precocial chicks. However, the MLC may be methodologically problematic for these pictures. Measuring mean luminance across the ROI may not capture the outline diffusion when both object and background are not monochromatic coloured but consist of a mottled pattern, which is frequently the case for natural habitats.

Altering the characteristics of appendages, background and predator vision had mechanism-specific consequences. As we concluded that reduction of edge intensity is the more likely mechanism, we restrict our discussion here to the impact of parameter changes on edge intensity. In the artificial setup, we found that an intermediate number of regular appendages helped to conceal the outline of the monochromatic object best. Further, we found that appendage thickness, transparency and length heterogeneity influenced outline concealment. They altered the optimal number of appendages needed and, in some cases, changed also the edge intensity. Protruding neoptile feathers of precocial chicks are thin, somewhat transparent and vary in the extent to which they stand out from the outline. Our results show that thicker appendages would lead overall to higher detectability and in that case, fewer appendages would lead to better concealment. In contrast, higher transparency required more appendages for best concealment. Similarly, we found that with increasing length heterogeneity more appendages were needed to achieve low edge intensities and reduce detectability.

Variation in spatial acuity is high across visual systems of different predators and had the largest effect on edge intensity. Intermediate to high appendage numbers reduced the edge intensity of the ROI most, regardless of spatial acuity of the simulated predator. Yet, mean edge intensities were highest for the simulated system with the highest spatial acuity. From the same viewing distance, predators with high spatial acuity, such as humans or birds of prey, perceive a lot more details of an object compared to predators with a lower spatial acuity such as canids or corvids (Caves et al. 2018). As spatial acuity decreases with viewing distance (Caves and Johnsen 2018), mammalian predators need to approach feathered chicks closer to detect their outline.

Interestingly, background complexity did not alter the optimal number of appendages nor impact overall edge intensities dramatically. Background complexity often makes detection of objects harder and therefore contributes to camouflage (Dimitrova and Merilaita 2010, Xiao and Cuthill 2016). The multicoloured fringed feathers themselves could contribute to increasing complexity. Such an effect would have the largest impact on a more uniform background. The mixture of appendages and background will also create new false edges and increase disruptive colouration (Troscianko et al. 2017). Nevertheless, any such effects by protruding feathers are likely to be small as the feather region is only very narrow and, hence, will only impact the immediate surrounding of the chick. Hence it is unclear whether this effect is biologically relevant for detection through predators.

One drawback of our study is that we did not test empirically whether the appendages indeed reduce detectability by predators, e.g. through a predation experiment (e.g. similar to (Cuthill et al. 2005, Farkas et al. 2013)). Measuring the detection time of objects with and without appendages similar to protruding neoptile feathers would be an important test for the relevance of this mechanism in nature. Concealing the outline is unlikely to be the main antipredator strategy of chicks. We rather suggest that it works in concert with the cryptic colouration of the downy plumage, chick behaviour such as finding optimal hiding places and predator distraction or defence through their parents. Yet our results regarding the spatial acuity suggest that the fringed feathers could be an important component of a visual antipredator strategy against mammalian predators. Even if the reduction in detectability is only small, concealing the outline may enhance survival of precocial chicks during early life when chicks face a very high predation risk (Colwell et al. 2007, Brudney et al. 2013, Eberhart-Phillips et al. 2018), especially as the costs for having the protruding feathers may not be high.

Appendages that alter the outline are commonly found in nature. Examples of vertebrates with irregular outlines are known, e.g. from cephalopods (Panetta et al. 2017), fish (Allen et al. 2015), amphibians (Rauhaus et al. 2012) and reptiles (Buxton 1923). A striking example is provided by many insect larvae such as hairy caterpillars which, as chicks, have typically reduced mobility in comparison with the adult form. Birds have a strong influence on caterpillar mortality (Campbell and Sloan 1977), but hairy caterpillars are less preferred prey for avian predators than non-hairy caterpillars (Whelan et al. 1989). We suggest that concealing the outline might be one currently underappreciated function of hairy appendages contributing to improved camouflage.

## Conclusion

The ‘irregular marginal form’ as a camouflage strategy has inspired early researchers on camouflage (Cott 1940) but evidence for this mechanism so far has been limited. Our results suggest that body appendages such as feathers or hairs can help to create an ‘irregular marginal form’ that serves to diffuse the object outline. Appendages with the characteristics of protruding neoptile feathers reduced the edge intensity in a proof of principle analysis and on images of precocial chicks taken in their natural environment. Appendages also served to reduce mean luminance differences when both object and background were uniformly coloured but this mechanism failed to contribute to outline diffusion when we analysed images of chicks in their natural backgrounds. Improved camouflage through outline diffusion could be an important function of heterogenous integuments which are found in a variety of organisms.

## Acknowledgements

We thank Salvador Del Angel Gómez, Medardo Cruz-López and Ivan Guardado González for help with fieldwork. We are grateful to Mary Caswell Stoddard and the members of the research group Behavioural Genetics and Evolutionary Ecology for discussion of methodology and results.

## Declarations

### Data availability statement

Raw images, data and script are stored in Edmond the Open Research Data Repository of the Max Planck Society (https://edmond.mpdl.mpg.de/imeji/).

### Author contributions

VAR, CK and DM conceptualised the study. TV carried out field work. VAR and CK analysed, interpreted the data and wrote the manuscript. All authors revised the manuscript.

Permits – Fieldwork permits to collect the data were granted by the Secretaría de Medio Ambiente y Recursos Naturales (SEMARNAT). All field activities were performed in accordance with the approved ethical guidelines outlined by SEMARNAT.

