## Supplementary Material for "Fluffy feathers: how neoptile feathers contribute to camouflage in precocial chicks"

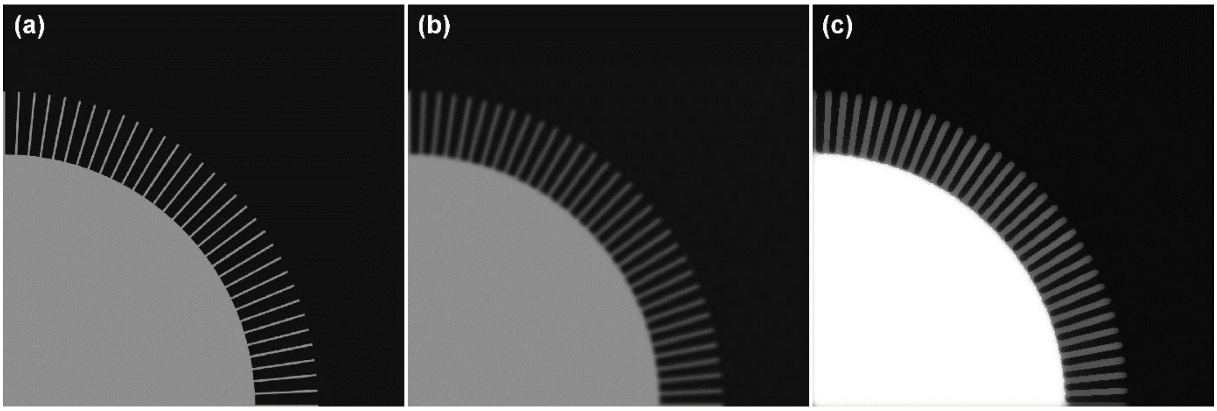


**Figure S1:** Modelling human vision with an acuity of 72 cpd. (a) Cone catch image (b) Gaussian Acuity Control (c) Edge reconstruction with Receptor Noise Limited (RNL) filter. All images exist as a stack of 4 channels (long, medium and short wave, and luminance).

**Table S1:** Parameter combinations used in Experiment 1. Differences to Basic Scenario indicated in shaded cells

| Parameter comb. index | Scenario | Number of appendages | Appendage thickness (Pt/px/mm) | Distance between appendages (px/mm) | Appendage transparency | Length hetero-geneity | Background | Acuity (cpd) |
|---|---|---|---|---|---|---|---|---|
| a | Basic Scenario | 0 | 1/4/0.4 | - | 0% | all 100% length | dark grey | 72 |
| b | | 32 | | 88/7.5 | | | | |
| c | | 64 | | 42/3.6 | | | | |
| d | | 128 | | 19/1.6 | | | | |
| e | | 256 | | 7/0.6 | | | | |
| f | | 512 | | 2/0.1 | | | | |
| g | | full circle | | - | | | | |
| h | Scenario 1a: 2 Pt Thickness | 0 | 2/8/0.7 | - | 0% | all 100% length | dark grey | 72 |
| i | | 32 | | 84/7.1 | | | | |
| j | | 64 | | 38/3.2 | | | | |
| k | | 128 | | 15/1.3 | | | | |
| l | | 256 | | 3/0.3 | | | | |
| m | | 512 | | -2/-0.2 | | | | |
| n | | full circle | | - | | | | |
| o | Scenario 1b: 3 Pt Thickness | 0 | 3/12/1.1 | - | 0% | all 100% length | dark grey | 72 |
| p | | 32 | | 80/6.8 | | | | |
| q | | 64 | | 34/2.9 | | | | |
| r | | 128 | | 11/0.9 | | | | |
| s | | 256 | | -1/-0.1 | | | | |
| t | | 512 | | -7/-0.6 | | | | |
| u | | full circle | | - | | | | |
| v | Scenario 2a: 25% Transparency | 0 | 1/4/0.4 | - | 25% | all 100% length | dark grey | 72 |
| w | | 32 | | 88/7.5 | | | | |
| x | | 64 | | 42/3.6 | | | | |
| y | | 128 | | 19/1.6 | | | | |
| z | | 256 | | 7/0.6 | | | | |
| aa | | 512 | | 2/0.1 | | | | |
| ab | | full circle | | - | | | | |
| ac | Scenario 2b: 50% Transparency | 0 | 1/4/0.4 | - | 50% | all 100% length | dark grey | 72 |
| ad | | 32 | | 88/7.5 | | | | |
| ae | | 64 | | 42/3.6 | | | | |
| af | | 128 | | 19/1.6 | | | | |
| ag | | 256 | | 7/0.6 | | | | |
| ah | | 512 | | 2/0.1 | | | | |
| ai | | full circle | | - | | | | |

| | | | | | | | | |
|---|---|---|---|---|---|---|---|---|
| aj | Scenario 3a: 1/2 at 50% length | 0 | 1/4/0.4 | - | 0% | 1/2 at 50% length | dark grey | 72 |
| ak | | 32 | | 88/7.5 | | | | |
| al | | 64 | | 42/3.6 | | | | |
| am | | 128 | | 19/1.6 | | | | |
| an | | 256 | | 7/0.6 | | | | |
| ao | | 512 | | 2/0.1 | | | | |
| ap | | full circle | | - | | | | |
| aq | Scenario 3b: 1/2 at 25% & 1/4 at 50% length | 0 | 1/4/0.4 | - | 0% | 1/2 at 25% & 1/4 at 50% length | dark grey | 72 |
| ar | | 32 | | 88/7.5 | | | | |
| as | | 64 | | 42/3.6 | | | | |
| at | | 128 | | 19/1.6 | | | | |
| au | | 256 | | 7/0.6 | | | | |
| av | | 512 | | 2/0.1 | | | | |
| aw | | full circle | | - | | | | |
| ax | Scenario 4a: background - big tiles | 0 | 1/4/0.4 | - | 0% | all 100% length | chessboard fields 346 px/29.3 mm | 72 |
| ay | | 32 | | 88/7.5 | | | | |
| az | | 64 | | 42/3.6 | | | | |
| ba | | 128 | | 19/1.6 | | | | |
| bb | | 256 | | 7/0.6 | | | | |
| bc | | 512 | | 2/0.1 | | | | |
| bd | | full circle | | - | | | | |
| be | Scenario 4b: background - small tiles | 0 | 1/4/0.4 | - | 0% | all 100% length | chessboard fields 86 px/7.3 mm | 72 |
| bf | | 32 | | 88/7.5 | | | | |
| bg | | 64 | | 42/3.6 | | | | |
| bh | | 128 | | 19/1.6 | | | | |
| bi | | 256 | | 7/0.6 | | | | |
| bj | | 512 | | 2/0.1 | | | | |
| bk | | full circle | | - | | | | |
| bl | Scenario 5a: 30 cpd acuity | 0 | 1/4/0.4 | - | 0% | all 100% length | dark grey | 30 |
| bm | | 32 | | 88/7.5 | | | | |
| bn | | 64 | | 42/3.6 | | | | |
| bo | | 128 | | 19/1.6 | | | | |
| bp | | 256 | | 7/0.6 | | | | |
| bq | | 512 | | 2/0.1 | | | | |
| br | | full circle | | - | | | | |
| bs | Scenario 5b: 10 cpd acuity | 0 | 1/4/0.4 | - | 0% | all 100% length | dark grey | 10 |
| bt | | 32 | | 88/7.5 | | | | |
| bu | | 64 | | 42/3.6 | | | | |
| bv | | 128 | | 19/1.6 | | | | |
| bw | | 256 | | 7/0.6 | | | | |
| bx | | 512 | | 2/0.1 | | | | |
| by | | full circle | | - | | | | |

**Table S2:** Spatial acuity of humans and potential chick predators.

| Spatial acuity approximation | Potential predator of snowy plover chicks | Related species with known spatial acuity | Spatial acuity |
|---|---|---|---|
| High (72 cpd) | Human *(Homo sapiens)* | human (*Homo sapiens*) – (Land 1981, Hirsch and Curcio 1989, Land and Nilsson 2012, Caves and Johnsen 2018) | 72 – 73 cpd |
| | Birds of prey (Page et al. 1985, Mabee and Estelle 2000) | Brown Falcon (*Falco berigora*) – (Reymond 1987) | 73 cpd |
| Medium (30 cpd) | Crested caracaras (*Caracara cheriway*) – (C Küpper, personal observations) | Chimango caracara (*Phalcoboenus chimango*) – (Potier et al. 2016, Caves et al. 2018) | 15 – 40 cpd |
| | Corvids – (Page et al. 1985, Mabee and Estelle 2000) | Several corvids – (Dabrowska 1975, Caves et al. 2018) | 30 – 33 cpd |
| | Racoon (*Procyon lotor*) – (Stoddard et al. 2016) | Racoons (*Procyon lotor*) – (Johnson and Michels 1958) | 25 – 30 cpd |
| Low (10 cpd) | Feral dog (*Canis familiaris*) – (Stoddard et al. 2016) | Dog (*Canis familiaris*) – (Bromberg and Dawson 1980, Odom et al. 1983, Pretterer et al. 2004) | 4.62 – 12.59 cpd |
| | Coyote (*Canis latrans*) – (Stoddard et al. 2016) | | |
| | Bobcat (*Lynx rufus*) – (Stoddard et al. 2016) | European lynx (*Lynx europea*) – (Maffei et al. 1990) | 7 – 8 cpd |
| | | Cat (*Felis catus*) – (Wässle 1971, Caves and Johnsen 2018) | 10 cpd |

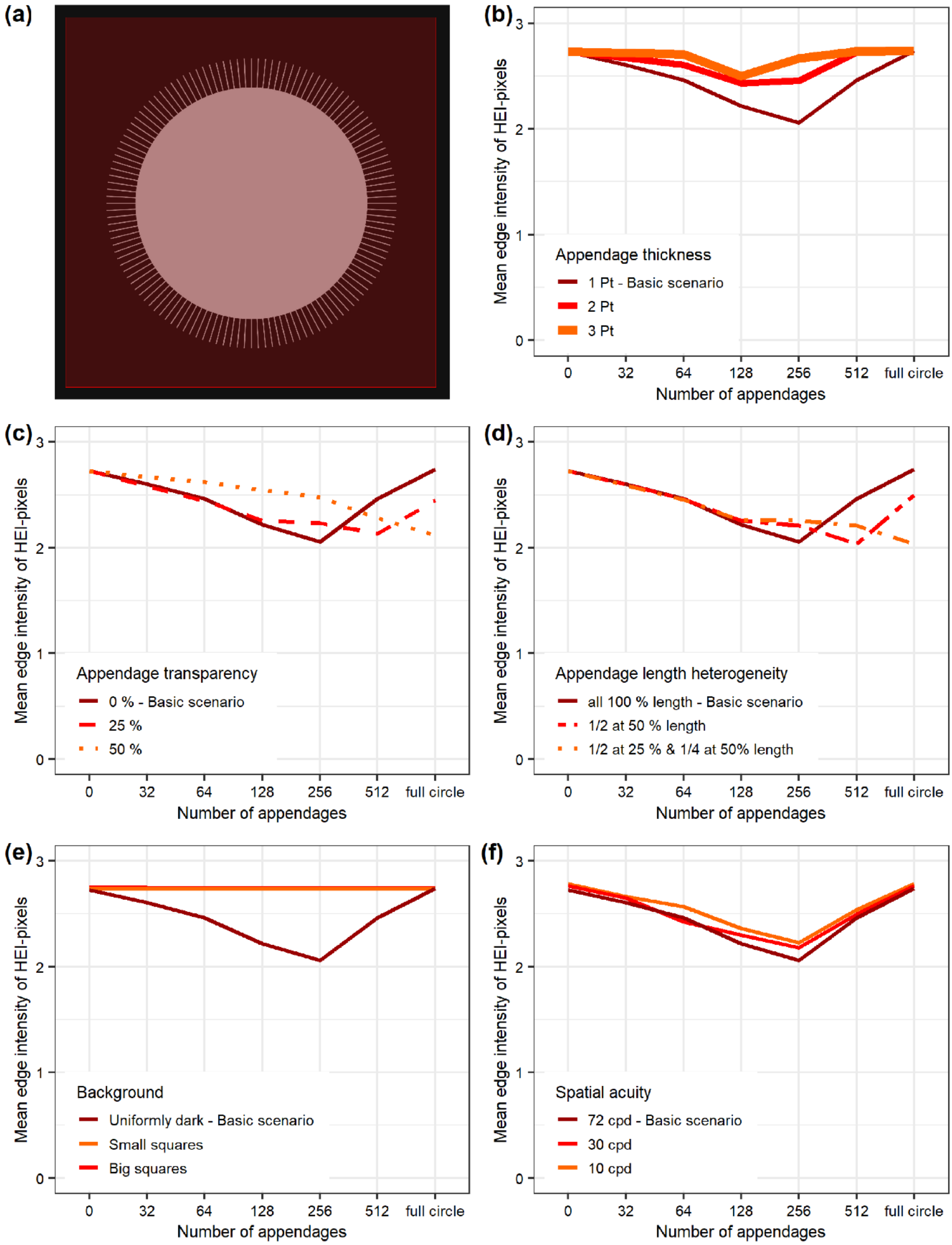


**Figure S2:** Local edge intensity analysis in experiment 1 with an expanded region of interest (ROI). The highest 0.37 % of the pixels are classified as "High Edge Intensity-pixels" (HEI-pixels). (a) ROI is a 1500 x 1500 pixel-wide rectangle (red). (b) Scenario 1: variation in appendage thickness. (c) Scenario 2: variation in appendage transparency. (d) Scenario 3: variation in appendage length. (e) Scenario 4: variation in background complexity. Note that the "Big squares" and "Small squares" curves overlap fully. (f) Scenario 5: variation in spatial acuity.

**Table S3:** General characterisation of the chick images. Percentage values of ROI available for analysis.

| Chick ID | Feather region excl. shadow | Contour region excl. shadow | HEI-pixels threshold |
|---|---|---|---|
| CN0333 | 77% | 75% | 0.9859 |
| CN0339 | 57% | 58% | 0.9827 |
| CN0340 | 61% | 62% | 0.9789 |
| CN0345 | 88% | 82% | 0.9815 |
| CN0347 | 61% | 59% | 0.9839 |
| CN0350 | 82% | 82% | 0.9829 |
| CN0353 | 91% | 89% | 0.9789 |
| CN0356 | 58% | 58% | 0.9830 |
| CN0360 | 64% | 64% | 0.9854 |
| CN0361 | 100% | 100% | 0.9840 |
| CN0363 | 100% | 100% | 0.9789 |
| CN0364 | 81% | 78% | 0.9846 |
| CN0367 | 58% | 56% | 0.9858 |
| CN0411 | 66% | 56% | 0.9821 |
| CN0415 | 57% | 55% | 0.9800 |
| Mean | 73% | 72% | 0.9826 |
